# Dynamic and atypical E-cadherin-based attachments mediate melanoblast migration through confined epithelial spaces

**DOI:** 10.1101/2025.08.20.671360

**Authors:** Denay J.K. Richards, Brandon M. Trejo, Parijat Sil, Abhishek Biswas, Rebecca A. Jones, Lionel Larue, Danelle Devenport

## Abstract

Epithelial tissues are populated with accessory cells such as immune cells, sensory cells, and pigment-producing melanocytes, which must migrate through and intercalate between tightly adherent epithelial cells. Although much is known about how cells migrate through interstitial spaces consisting of predominantly of collagen-rich ECM and mesenchyme, how cells migrate through confined epithelial spaces without impairing barrier function is far less understood. Here, using live imaging of the mouse epidermis, we captured the migration of embryonic melanocytes (melanoblasts) while simultaneously visualizing the basement membrane or epithelial surfaces. We show that melanoblasts migrate through both basal and suprabasal layers of the epidermis and hair follicles where they use keratinocyte surfaces, as well as the basement membrane, as substrates for migration. We show that melanoblasts form atypical and dynamic E-cadherin based attachments to surrounding keratinocytes that largely lack the cytoplasmic catenins known to anchor E-cadherin to the actin cytoskeleton. We show E-cadherin is needed in both melanoblasts and keratinocytes to stabilize migratory protrusions, and that depleting E-cadherin in melanoblasts results in reduced motility and ventral depigmentation in adult mice. These findings illustrate how migratory cells co-opt the cell-cell adhesion machinery connecting adjacent epithelial cells to invade between and migrate through them without interrupting the skin barrier.

## INTRODUCTION

The epithelial layers of internal and external organs are interspersed with a multitude of accessory cells that protect from pathogens, detect sensory inputs and provide UV protection. These highly migratory cell types often travel long distances from their place of origin to target and colonize their resident tissues. Upon arrival they must infiltrate and squeeze between the epithelial layers held tightly together by cell-cell adhesions. Although much is known about the mechanisms by which cells migrate on 2D extracellular substrates or through 3D interstitial matrices (Alonso-Matilla et al., 2025; Pawluchin and Galic, 2022; SenGupta et al., 2021; Yamada and Sixt, 2019), how cells move between tightly adherent epithelial cells without impairing the epithelium’s barrier function is poorly understood. The skin epidermis, in which immune cells, sensory nerve endings and pigment producing melanocytes are interspersed among the skin’s many layers of stratified keratinocytes (Cui and Man, 2023; Heath and Carbone, 2013; Stander and Schmelz, 2024), is an excellent model organ to investigate how migratory accessory cells move through confined epithelial spaces.

Melanocytes are melanin-producing cells that reside in the epidermis and hair follicles and are responsible for pigmentation of the skin and hair (Cui and Man, 2023). Melanocytes form elaborate cellular protrusions that enable them to intercalate between and transfer melanin to surrounding epithelial cells called keratinocytes (Brombin and Patton, 2024; Mort et al., 2015). It is essential that melanocytes are dispersed throughout the entire skin surface to protect keratinocytes and other epidermal cells from UV radiation (UVR); failure of melanocyte precursors, known as melanoblasts, to effectively colonize the epidermis during development leaves regions of the skin susceptible to UVR-mediated DNA damage and at increased risk for a variety of skin cancers (Brombin and Patton, 2024; Cui and Man, 2023; Mort et al., 2015; Wang et al., 2009). Thus, it is important to understand how highly migratory melanoblasts transit through the epithelial layers of the skin to populate the epidermis.

Melanoblasts derive from the neural crest, multipotent progenitors that delaminate from the dorsal-most region of the neural tube (Brombin and Patton, 2024; Vandamme and Berx, 2019). From their dorsal origin, melanoblasts must migrate long-distances through the dermis to reach the ventral-most regions of the skin and extremities. Melanoblasts cross into the epidermis where they continue to migrate and proliferate to populate the skin surface and hair follicles (Cui and Man, 2023). Compared to the ECM- and fibroblast-rich dermis they initially traverse, melanoblasts encounter a completely distinct microenvironment in the epidermis. Layers of tightly packed epidermal keratinocytes adhere to one another via desmosomes, adherens junctions and tight junctions (Brandner et al., 2015; Johnson et al., 2014; Niessen, 2007; Sumigray and Lechler, 2015), through which melanoblasts must somehow intercalate without damaging epithelial integrity. Thus, migration within and between the dermal and epidermal compartments likely involve distinct molecular processes. It is known that, despite their non-epithelial origin, melanoblasts in the epidermis upregulate the adhesion proteins E-cadherin and P-cadherin (Jouneau et al., 2000; Nishimura et al., 1999), suggesting they may attach to keratinocytes via homotypic E-cadherin binding (Li et al., 2011). However, E-cadherin mediated adhesion is generally thought to restrict cell migration or to connect cells so they move as collectives (Friedl and Mayor, 2017; Gupta and Yap, 2021; Kourtidis et al., 2017; Mayor and Etienne-Manneville, 2016; Messer and McDonald, 2023; Mishra et al., 2019; Rubtsova et al., 2022). Yet melanoblasts move through the epidermis rapidly and individually, and upon contact, do not form homotypic attachments with other melanoblasts (Laurent-Gengoux et al., 2018; Mort et al., 2010; Mort et al., 2016). We hypothesize that melanoblasts form specialized and atypical E-cadherin based attachments to surrounding keratinocytes, enabling them to invade between the epidermal layers and to use the keratinocyte surface as a substrate for migration.

Here, we combine live and high-resolution imaging with conditional knockouts in the mouse embryonic skin to define the composition and function of E-cadherin-based adhesions between melanoblasts and keratinocytes. We discover that melanoblasts form extensive, atypical E-cadherin based attachments with surrounding keratinocytes in both basal and suprabasal layers of the epidermis. However, unlike keratinocyte-keratinocyte adherens junctions (AJs) that are enriched in cytoplasmic catenins, melanoblast AJs largely lack p120-, α- and β-catenin, cytoplasmic components of adherens junctions that strengthen adhesions through interactions with the actin cytoskeleton. We observe that as melanoblasts migrate through the epidermis, they extend multiple dendritic protrusions and exhibit elongated cell bodies as they squeeze through adjacent keratinocytes. Depletion of E-cadherin in either melanoblasts or keratinocytes, via selective Cre-recombinase mediated knockout, results in decreased dendritic protrusions, increased cell body circularity and impaired melanoblast migration. Using live imaging, we find this morphological change represents a reduction in the stability of migratory protrusions resulting in decreased melanoblast velocity and distance covered. This defect ultimately results in ventral depigmentation in adult mice, indicative of impaired melanoblast migration. Collectively, our data suggest that migratory melanoblasts use E-cadherin homotypic binding to interact with the surface of keratinocytes to stabilize filopodia and generate traction needed to facilitate directed migration in a tightly adherent epithelial environment.

## RESULTS

### Melanoblasts migrate along the surfaces of keratinocytes in both basal and suprabasal layers of the epidermis

To understand how migratory melanoblasts transit through tightly adherent epidermal cells, we performed live imaging of embryonic skin explants in which melanoblasts are labeled with membrane-GFP driven by *Tyrosinase-Cre* (*Tyr-Cre*) and keratinocytes are labeled with membrane-tdTomato (*Tyr-Cre; mTmG*)(Cetera et al., 2018; Delmas et al., 2003; Muzumdar et al., 2007)(Figure 1A, Movie S1). This allowed us to simultaneously visualize the epithelial environment as melanoblasts migrate between keratinocytes to colonize the epidermis. Melanoblasts reside within the basal layer of the epidermis and are thought to migrate upon the basement membrane (BM), a sheet of specialized extracellular matrix (ECM) separating the epidermis from the underlying dermis (Figure 1B)(Haage et al., 2020; Haass et al., 2005; Li et al., 2011). In confocal z-stacks through the epidermis, we observed melanoblasts migrating between keratinocytes in the basal layer, but found they were also prevalent within the suprabasal layers (Figure 1C-D). Suprabasal melanoblasts were most often observed migrating above and through the center of developing hair follicles (Figure 1E, Movie S2), suggesting melanoblasts can use keratinocyte surfaces as substrates for migration.

**Figure 1.**
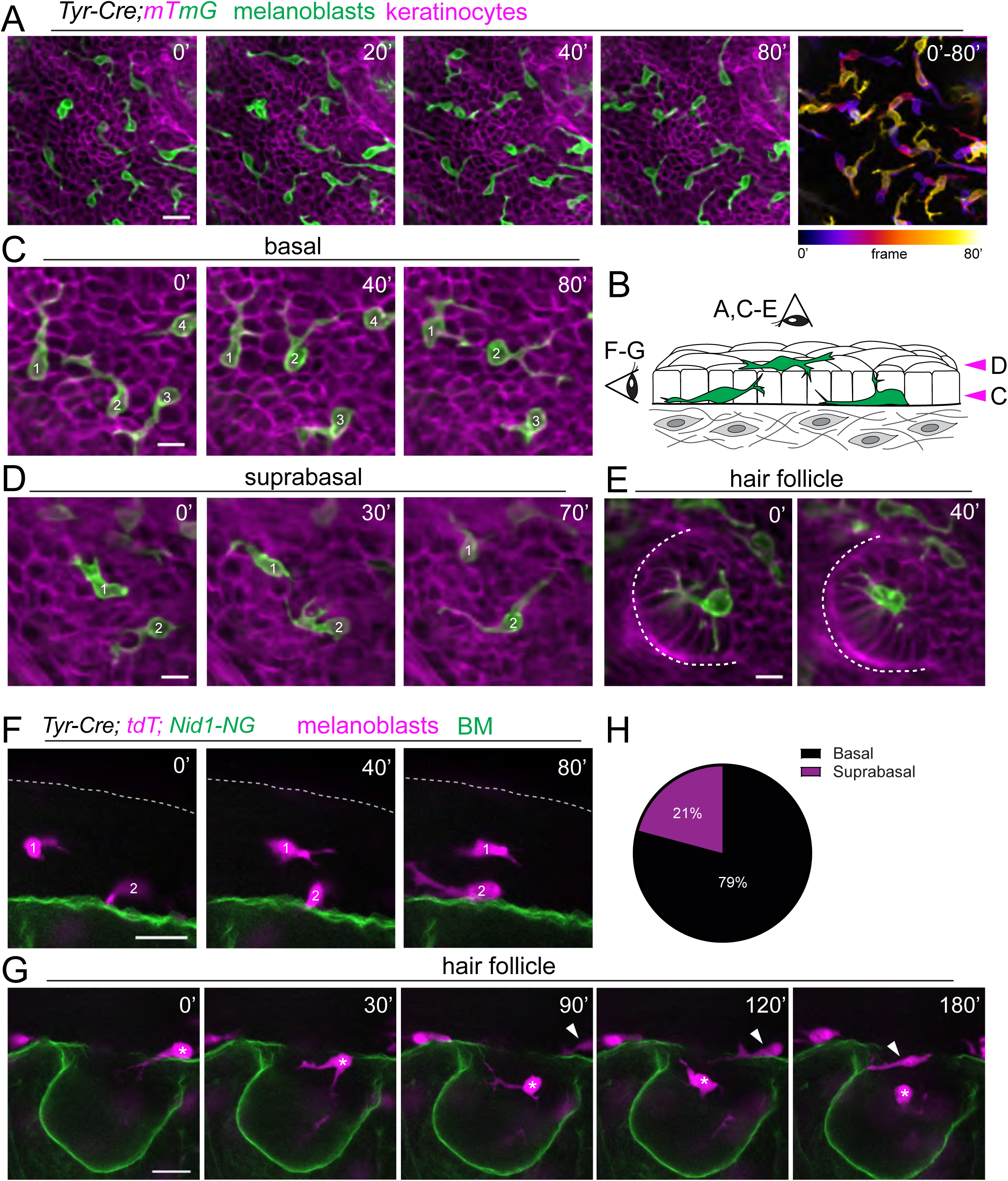
Melanoblasts migrate along the surfaces of keratinocytes in both basal and suprabasal layers of the epidermis. **(A)** Still frames from live imaging of epidermal explants from E16.5 *Tyr-Cre; mTmG* embryos. Melanoblasts (green, mG) migrate between keratinocytes (magenta, mT). Right hand panel shows merged time frames to illustrate the extent of movement. Scale bar, 20um **(B)** Schematic representation of E16.5 epidermis in sagittal view. Arrowheads represent planes of imaging in C-D. **(C-E)** Single optical slices from confocal z-stacks are shown through the basal (A) and suprabasal (B) layers of the epidermis, and developing hair placode (E). Melanoblasts (green, mG) migrate between keratinocytes (magenta, mT) in the basal layer (C) and the suprabasal layers of the epidermis (D) and hair placodes (E). Scale bars, 10um. (**F**) Still frames from PS Multiview imaging of E16.5 *Tyr-Cre; tdTomato; Nid1-NG* whole mount explants. Melanoblasts (magenta, tdT) migrate both on the basement membrane (green, NG) and within the suprabasal layers of the skin. Scale bar, 20um (**G**) Still frames from PS Multiview imaging of skin explant from E16.5 *Tyr-Cre; tdTomato; Nid1-NG* embryo shows E16.5 melanoblasts migrating within the inner, suprabasal layers of a developing hair follicle and making few connections to the underlying basement membrane. Scale bar, 20um **(H)** Quantification of melanoblast cell bodies located in the suprabasal and basal layers in E16.5 *Tyr-Cre; tdTomato; Nid1-NG* whole mount explants.

To verify that melanoblasts can migrate between keratinocyte layers without appreciable contact with the ECM, we crossed *Tyr-Cre;tdTomato* mice to a newly-developed BM reporter mouse line in which endogenous Nidogen 1 is tagged with mNeonGreen (*Nid1-NG*)(Madisen et al., 2010)(Richards, in preparation). Using PS Multiview imaging to visualize the tissue in both planar and sagittal (PS) views (Jones et al., 2024), we confirmed that although melanoblasts do reside in close proximity to the basement membrane, many cells migrate above the basal layer, losing most if not all of their contacts to the ECM (Figure 1F, Movie S3). Quantification of basal versus suprabasal melanoblasts in scanning confocal z-stacks of fixed whole mount epidermis revealed that 20% of melanoblast cell bodies were located in the suprabasal layers (Figure 1H). Moreover, sagittal views from PS Multiview imaging of developing hair follicles showed that melanoblasts also migrate through the suprabasal cells located in the center of developing hair follicles, far from the nearest BM (Figure 1G, Movie S4). These data indicate that, in addition the ECM, melanoblasts use keratinocyte surfaces as substrates for migration.

### Melanoblasts form extensive E-cadherin based attachments to keratinocytes

Epithelial cells form cell-to-cell attachments via homotypic binding of classical cadherins. Melanoblasts are known to dramatically upregulate E-cadherin expression around the time they cross into the epidermis (Jouneau et al., 2000; Nishimura et al., 1999), and we hypothesized that melanoblasts use E-cadherin not only to attach to keratinocytes but also to invade between them and perhaps to generate traction forces for migration. To begin to test this hypothesis, we first performed a quantitative analysis of cadherin expression and localization in E16.5 *Tyr-Cre; mTmG* embryonic skins labeled with antibodies against E- or P-cadherin. The two-color membrane reporter allowed us to unambiguously distinguish melanoblast-keratinocyte (MB-KC) interfaces from those between neighboring keratinocytes (KC-KC). In the interfollicular epidermis, E-cadherin localized to both KC-KC and MB-KC interfaces and was present along melanoblast cell bodies (Li et al., 2011) as well as their dendritic protrusions (Figure 2A). P-cadherin, although upregulated in developing hair follicles, was relatively low in interfollicular keratinocytes and was largely absent from MB-KC junctions of interfollicular melanoblasts (Figure 2B)(Hardy and Vielkind, 1996; Muller-Rover et al., 1999). To measure cadherin levels, cell edges of keratinocytes and melanoblasts were segmented using the mTmG membrane labels and the mean fluorescence intensity of E- or P-cadherin was calculated per cell (Leybova et al., 2024). Whereas both interfollicular melanocytes and keratinocytes express low levels of P-cadherin, we found that melanoblasts express high levels of E-cadherin, comparable to or even higher than that of surrounding keratinocytes (Figure 2C). We conclude from these data that E-cadherin is the primary classical cadherin expressed by epidermal melanoblasts and that, despite being highly migratory, melanoblasts form E-cadherin-based attachments at levels comparable to or even greater than those between non-motile keratinocytes.

**Figure 2.**
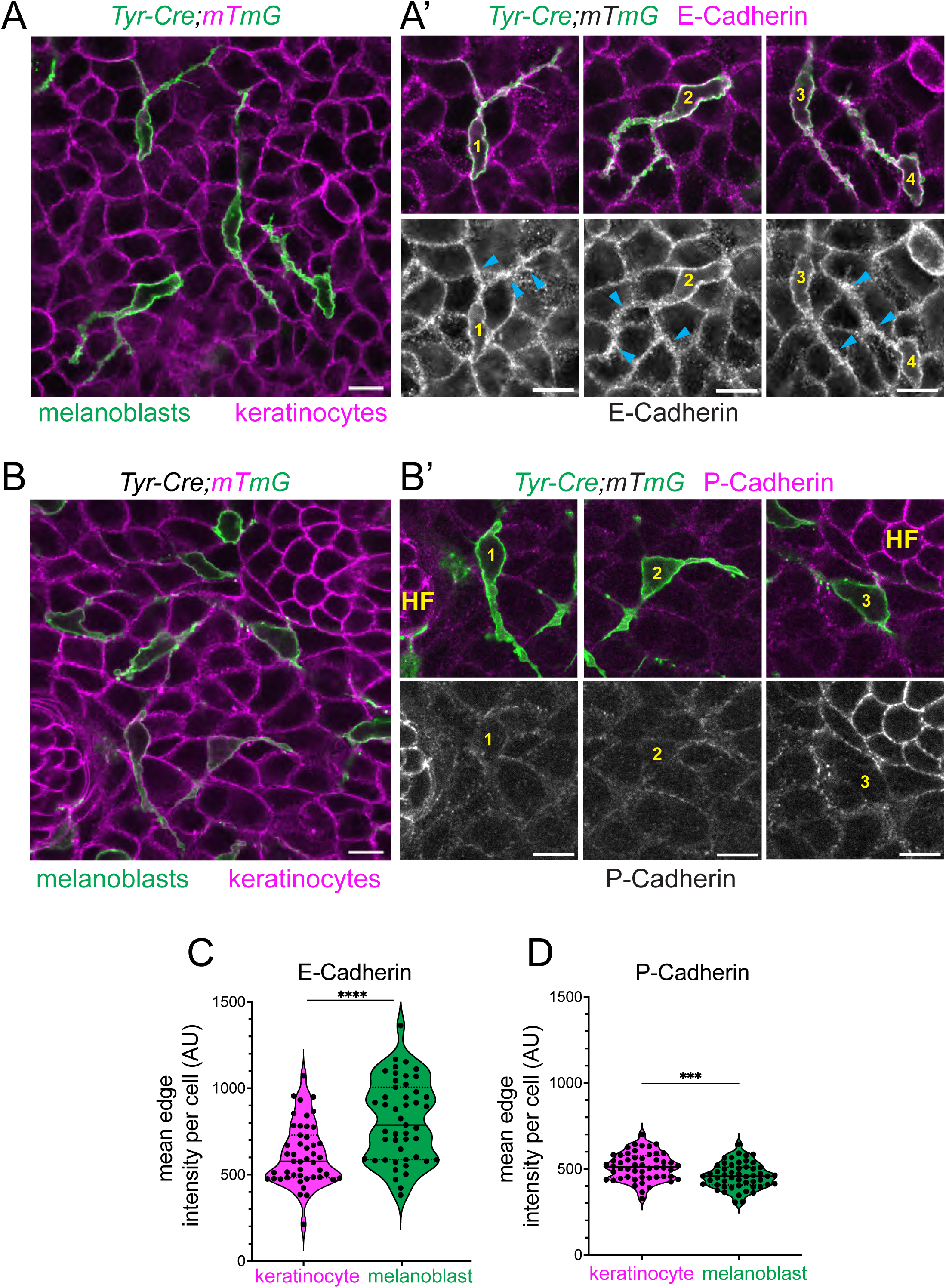
Melanoblasts form primarily E-cadherin-based attachments to interfollicular keratinocytes. (**A-B**) Representative planar views of wholemount skin explants from E16.5 *Tyr-Cre; mTmG* embryos labeled with E-cadherin (A) or P-cadherin (B). In A and B, melanoblast membranes are labeled green (mG) and keratinocyte membranes are magenta (mT). In A’ and B’ zoomed in views of individual melanoblasts are shown (numbered in A-B), with E-cadherin or P-cadherin shown in magenta (top panels) and in grayscale (bottom panels). Note that E-cadherin localizes to MB-KC cell interfaces, whereas P-cadherin does not. HF= hair follicle. Scale bars, 10um. (**C**) Quantification of mean E-cadherin intensity along MB-KC and KC-KC interfaces. Each point is the mean edge intensity per cell. n= 47 melanoblasts and 47 keratinocytes across three E16.5 skins. Two-tailed, unpaired t-test p< 0.0001. (**D**) Quantification of mean P-cadherin intensity along MB-KC and KC-KC interfaces. Each point is the mean edge intensity per cell. n= 52 melanoblasts and 52 keratinocytes across three E16.5 skins. Two-tailed, unpaired t-test p=0.0005.

### Conditional depletion of melanoblast E-cadherin reduces dendritic complexity and protrusion stability

As melanoblasts migrate through the epidermis, they form long migratory protrusions that extend in multiple directions (Haage et al., 2020; Li et al., 2011; Ma et al., 2013; Woodham et al., 2017). Long protrusions can further branch into smaller secondary and tertiary filopodia that extend and retract, sensing their local environment. To undergo forward directed motion, melanoblasts polarize by stabilizing a single primary protrusion and retracting others and then move in the direction of its main, stable projection, or pseudopod (Figure 1A, Movie S1). Melanoblasts frequently change direction through the extension and stabilization of new pseudopods (Laurent-Gengoux et al., 2018; Li et al., 2011; Mort et al., 2016). To understand how E-cadherin-mediated adhesion between melanoblasts and keratinocytes may contribute to the elongated and protrusive cellular morphology that accompanies melanoblast migration, we generated mouse embryos in which E-cadherin was conditionally deleted in melanoblasts using *Tyr-Cre* and quantified various morphological features of melanoblasts with and without E-cadherin (Boussadia et al., 2002). To validate that E-cadherin was efficiently depleted from melanoblasts using *Tyr-Cre*, we performed E-cadherin antibody labeling in whole mount skin explants from control and *Tyr-Cre; E-Cad fl/fl; mTmG* and embryos. Compared to control melanocytes, which express high levels of E-cadherin localized to interfaces with surrounding keratinocytes, *Tyr-Cre; E-Cad fl/fl; mTmG* (MB E-Cad cKO) melanoblasts lacked E-cadherin expression (Figure S1A-B). Moreover, compensatory P-cadherin expression was not upregulated in melanoblasts lacking E-cadherin, as is observed when E-cadherin is depleted in epidermal keratinocytes (Figure S1C)(Leybova et al., 2024; Tinkle et al., 2004; Tinkle et al., 2008). To compare the morphology of melanoblasts expressing or lacking E-cadherin, we acquired confocal z-stacks through the epidermal layers of control and *Tyr-Cre; E-Cad fl/fl; mTmG* whole mount skin explants and generated z-projections of the membrane-GFP channel. This allowed us to visualize in 2D the complex morphologies of melanoblast cell bodies and their dendritic projections which extend between epidermal layers in 3-dimensions (Figure 3A and D). We observed that melanoblasts lacking E-cadherin display fewer primary dendritic protrusions extending from their cell bodies than their wild-type counterparts (Figure 3B, E and G). Further, the protrusions that formed in E-Cad cKO melanoblasts were shorter and had fewer branch points (Figure 3C, F, H, I and Figure S1D). We measured the circularity of melanoblast cell bodies and found that those lacking E-cadherin were less elongated and significantly more circular than controls (Figure 3A, D, J and Figure S1C).

**Figure 3.**
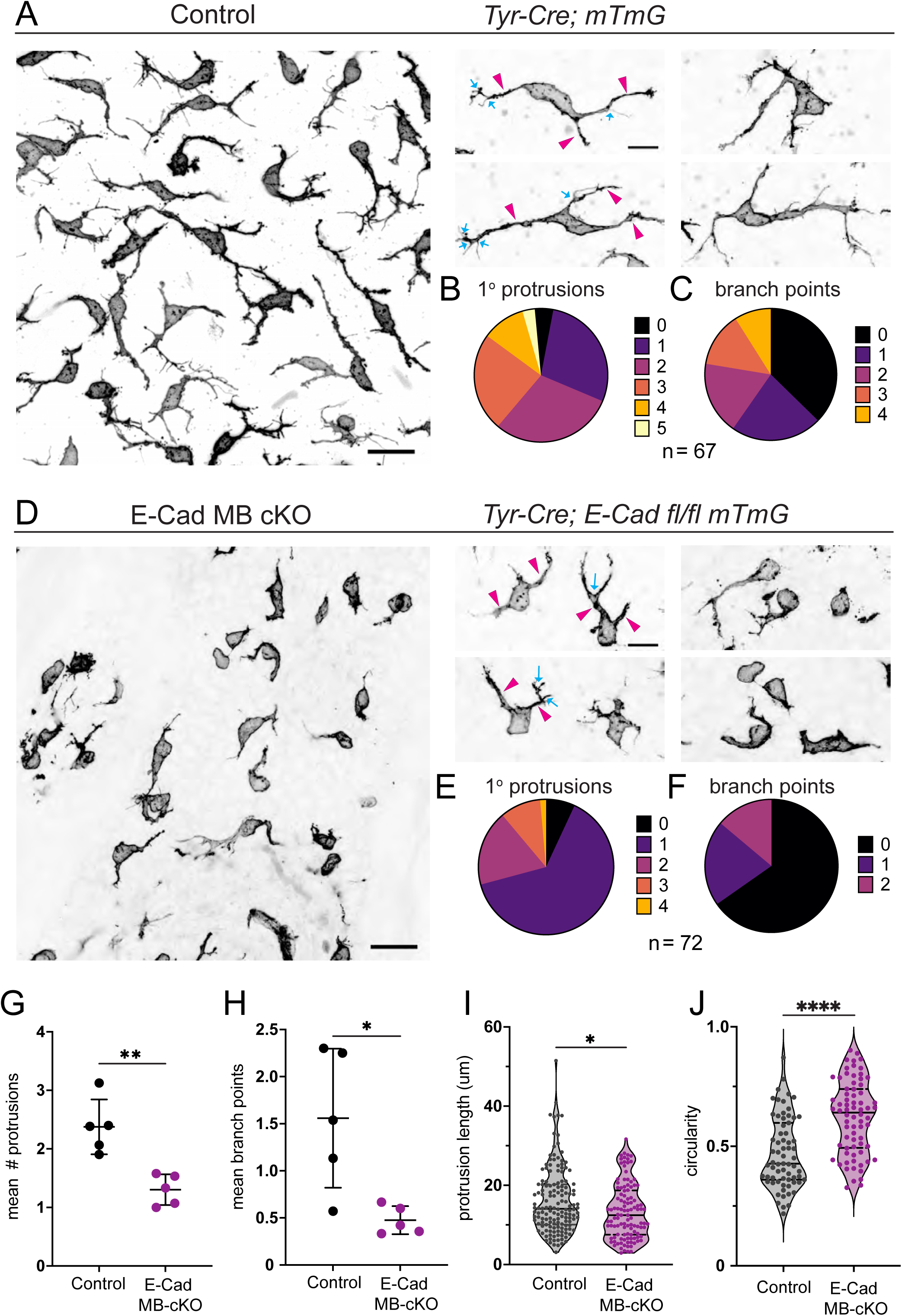
Melanoblast E-cadherin is required for dendritic complexity and pseudopod stability. Representative planar view of wholemount skin explant from E16.5 *Tyr-Cre; mTmG* embryo. Shown is a z-projection of the mG channel labeling melanoblasts in inverted grayscale to highlight melanoblast morphology. Scale bar, 20um. Right panels show zoomed in views of representative control melanoblasts. Arrowheads point to primary protrusions, blue arrows point to branch points. Scale bar, 10um. **(B-C)** Pie charts depicting the number of primary protrusions (B) and branch points per primary protrusion (C) displayed by control melanoblasts. n=67 MBs across five embryos. **(D)** Representative planar view of wholemount skin explant from E16.5 *Tyr-Cre; E-Cad fl/fl; mTmG* embryo. Z-projection of the mG channel labeling melanoblasts in inverted grayscale is shown. Scale bar, 20um. Right panels show zoomed in views of representative E-Cad cKO melanoblasts. Arrowheads point to primary protrusions, blue arrows point to branch points. Scale bar, 10um. **(E-F)** Pie charts depicting the number of primary protrusions (E) and branch points (F) displayed by E-Cad cKO melanoblasts. n=72 MBs across five embryos. **(G)** Mean number of primary protrusions across melanoblasts per biological replicate. n=5 control and n=5 E-Cad MB cKO backskins. Mann-Whitney test p=0.0079. **(H)** Mean branch points across melanoblasts per biological replicate. n=5 control and n=5 E-Cad MB cKO backskins. Mann-Whitney test p=0.0317. **(I)** Length of primary protrusions in um. Mann-Whitney test p=0.0113 **(J)** Circularity ratio. Mann-Whitney test p>0.0001. For G-J, n=67 melanoblasts across n=5 control embryos, and n=72 melanoblasts across n=5 E-Cad MB cKO embryos. Mean +/- S.D. are shown.

To determine whether the reduced protrusivity observed in E-cadherin cKO melanoblasts is due to an inability to form pseudopodial extensions or to stabilize them, we performed time-lapse imaging of control (*Tyr-Cre; E-Cad +/+; mTmG*) and E-Cad MB cKO (*Tyr-Cre; E-Cad fl/fl; mTmG*) skin explants to visualize protrusion dynamics of actively migrating melanoblasts in live skin (Figure 4A-C, Movie S5 and S6). Melanoblasts extended both long and short protrusions that often branched or bifurcated along their length (Figure 4A, Movie S5). The tips of protrusions were particularly dynamic, extending and retracting numerous filopodia (Figure 4C, Movie S5). In contrast to the reduced number of primary protrusions observed in fixed samples, we found that live E-cadherin cKO melanoblasts were able to extend both long and short protrusions (Figure 4B, Movie S6).

**Figure 4.**
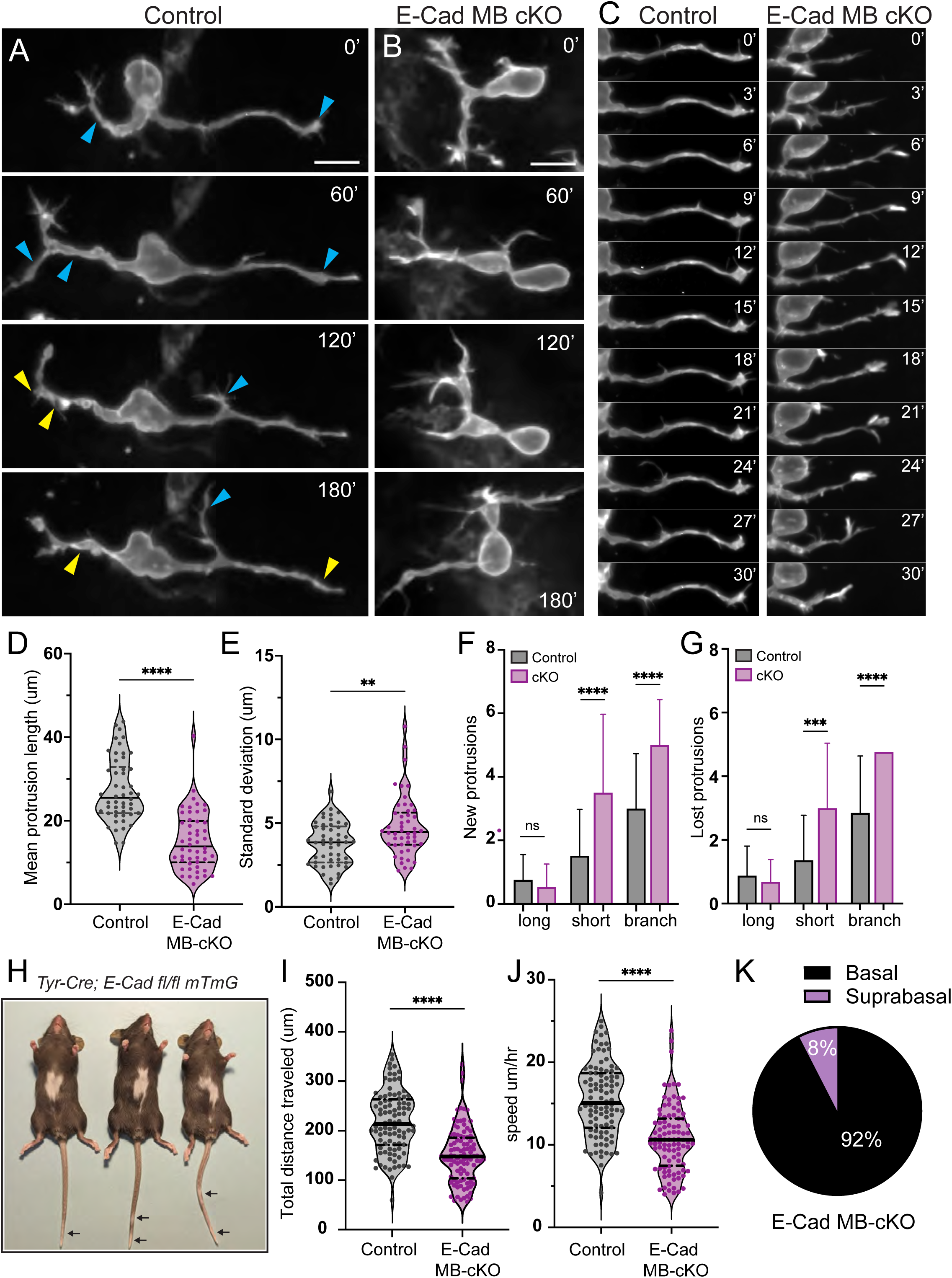
Defects in protrusion stability and migration resulting from depleting E-cadherin in melanoblasts. **(A)** Time-lapse still frames from live imaging of E16.5 *Tyr-Cre; mTmG* epidermal explants. Membrane-GFP channel of representative control melanoblast is shown. Images were captured every 3 minutes for 3 hours to capture protrusion dynamics (see Supplemental Movie 5). Blue arrowheads indicate extending protrusions, yellow arrowheads point to retracting protrusions. Scale bar, 10um. **(B)** Still frames from time-lapse imaging of E16.5 *Tyr-Cre; E-Cadfl/fl; mTmG* epidermal explants. Membrane-GFP channel from representative E-Cad MB cKO melanoblast is shown (see Supplemental Movie 6). Scale bar, 10um. **(C)** Still frames of representative protrusions from *Tyr-Cre; mTmG* (control) and *Tyr-Cre; E-Cadfl/fl; mTmG* (Ecad MB cKO) melanoblasts captured every 3 minutes for 30 minutes. (see Supplemental Movie 7). **(D)** Quantification of mean protrusion length. Each dot represents one primary protrusion, whose length was measured every 3 minutes for 30 minutes. n=50 melanoblast protrusions from 3 different embryos for each genotype. Significance was determined using two-tailed, unpaired t-test. p<0.0001. **(E)** Standard deviation of protrusion length. Each dot represents one primary protrusion, whose length was measured at 3-minute intervals for 30 minutes. n=50 melanoblast protrusions from 3 different embryos for each genotype. Significance was determined using two-tailed, unpaired t-test with Welch’s correction p=0.0014. **(F)** Number of new protrusions per melanoblast formed over 3 hour time-lapse. Control; n=33 melanoblasts from 3 embryos. E-Cad MB cKO; n=38 melanoblasts from 3 embryos. Two-tailed, unpaired t-tests: long p=0.2034; short p<0.0001, branch p=<0.0001 **(G)** Number of protrusions lost per melanoblast over 3 hour time-lapse. Control; n=33 melanoblasts from 3 embryos. E-Cad MB cKO; n=38 melanoblasts from 3 embryos. Two-tailed, unpaired t-tests: long p=0.3186; short p=0.0002; branch p<0.0001. **(H)** Three adult stage *Tyr-Cre; E-Cadfl/fl; mTmG* mice with white belly spotting phenotype. Arrows point to depigmented regions of the tail epidermis. **(I-J)** Quantification of melanoblast migration. Images were captured every 10 minutes for 14 hours and melanoblasts tracked to capture longer-term migration behaviors. (I) Total distance traveled (microns). Control; n=94 melanoblasts from 3 embryos. E-Cad MB cKO; n=96 melanoblasts from 3 embryos. Mann-Whitney test: p<0.0001. (J) Mean speed (um/hr). Control; n=94 melanoblasts from 3 embryos. E-Cad MB cKO; n=96 melanoblasts from 3 embryos. Mann-Whitney test: p<0.0001. **(K)** Proportion of melanoblast cell bodies located in the basal and suprabasal layers of the epidermis in E16.5 *Tyr-Cre; E-Cadfl/fl; mTmG* skins. n=53 melanoblasts across 3 embryos.

However, these protrusions were less stable than those of wild-type melanoblasts. Protrusions lengths over a 30-minute time period were significantly more variable in E-Cad MB cKO melanoblasts as they extended and retracted more frequently, resulting in shorter average protrusion lengths (Figure 4C-E, Movie S7). Additionally, melanoblasts lacking E-cadherin formed and lost a greater number of short and branched protrusions than controls (Figure 4F-G). The discrepancy in protrusion number and morphology between live and fixed tissue samples was not unexpected as filipodia-like extensions are known to be sensitive to fixation, and appear to be more sensitive in melanoblasts lacking E-cadherin. Thus, these live imaging experiments revealed that the reduced dendritic complexity observed in fixed E-Cad MB cKO explants is due to a significant decrease in protrusion stability rather than formation. Collectively, these data indicate that melanoblast E-cadherin is needed to stabilize migratory protrusions that extend between keratinocytes.

### Melanoblasts require E-cadherin for efficient migration and epidermal colonization

Melanoblasts migrate great distances from their origins in the dorsal neural crest to populate the skin surface (Cui and Man, 2023; Petit and Larue, 2016; Vandamme and Berx, 2019). Defects in melanoblast migration during embryonic development lead to epidermal depigmentation in adult stages, particularly in regions furthest from their origin such as the ventral trunk, and extremities of the limbs and tail (Crawford et al., 2020; Haage et al., 2020; Li et al., 2011; Ma et al., 2013; Woodham et al., 2017). To determine the global impact of selective E-cadherin knockout in melanoblasts, we allowed *Tyr-Cre; E-Cad fl/fl; mTmG* mice and their littermates to develop into adulthood. We found that *Tyr-Cre; E-Cad fl/fl; mTmG*, but not *Tyr-Cre; E-Cad fl/+; mTmG* adult mice displayed ventral coat depigmentation (Figure 4H), termed “white belly spotting”, a phenotype characteristic of defective melanoblast migration, survival and/or proliferation (Cable et al., 1995; Mackenzie et al., 1997; Mort et al., 2016; Steel et al., 1992; Wehrle-Haller and Weston, 1995). This phenotype in adults strongly suggested that the defects in protrusion dynamics observed in E-cadherin cKO melanoblasts result in an overall impairment their migration over longer distances through the skin. To test this, we performed time-lapse imaging of control (*Tyr-Cre; mTmG*) and E-Cad MB cKO (*Tyr-Cre; E-Cad fl/fl; mTmG*) E16.5 skin explants over a 14-hour time period (Movie S8). Using semi-automated cell tracking to chart their migration paths, we measured the distance, speed and persistence of migration. On average, E-cadherin cKO melanoblasts migrated more slowly and traveled a shorter total distance than wild type melanoblasts, but their persistence and straight line displacements were similar, and were indicative of non-directed and confined movement (Figure 4I-J, Movie S8). Additionally, melanoblasts lacking E-cadherin were located more frequently in the basal layer of *Tyr-Cre; E-Cad fl/fl; mTmG* skins, which was reflected as a decrease in the proportion of suprabasal melanoblasts compared to controls (8% vs 21% in controls; Figure 4K, compare to Figure 1H). These findings indicate that E-cadherin expression within melanoblasts is needed to generate persistent, directional migration through the epidermal microenvironment, and suggests the ventral depigmentation observed in adults results from a failure of embryonic melanoblasts lacking E-cadherin to complete migration to the ventral most regions. That a greater proportion of E-Cad cKO melanoblasts are located in the basal layer further suggests that melanoblasts require E-cadherin to attach to suprabasal cell surfaces for use as a substrate for migration. Without E-cadherin, melanoblasts rely on other substrates, likely the basement membrane ECM, present in the basal layer.

### E-cadherin is required non-autonomously in keratinocytes to stabilize melanoblast protrusions

Our data thus far demonstrate a cell autonomous requirement for E-cadherin within melanoblasts to generate stable protrusions and to sustain directional migration. One interpretation of these data is that homotypic E-cadherin attachments with neighboring keratinocytes function to stabilize migratory protrusions, and to aid in the generation of traction forces needed for forward motion. However, it is possible the loss of E-cadherin alters other aspects of melanoblast function that are secondary or unrelated to homotypic adhesion. Therefore, we asked whether E-cadherin expression within keratinocytes is required for wild-type melanoblasts to migrate.

This information is key to understanding whether the morphological and migratory changes seen in E-Cad MB cKO melanoblasts is due to disruption of MB-KC adhesion. To determine the impact of keratinocyte E-cadherin on wild-type melanoblast morphology, we generated *K14-Cre; E-Cad fl/fl; mTmG* (E-Cad KC cKO) embryos in which epidermal keratinocytes lack E-cadherin (Tinkle et al., 2004; Vasioukhin et al., 1999). In this model, K14 drives Cre expression in keratinocytes of the basal progenitor layer resulting E-cadherin deletion in all stratified layers. However, melanoblasts maintain E-cadherin expression and also express mTomato due to their lack of Cre expression driven by K14-Cre, allowing melanoblasts to be distinguished from mGFP-expressing keratinocytes (Figure 5A). Consistent with previous studies, we find that keratinocytes in the basal layer upregulate P-cadherin in the absence of epidermal E-cadherin (Figure 5B)(Leybova et al., 2024; Tinkle et al., 2004; Tinkle et al., 2008), and this is sufficient to maintain basal layer integrity (Tinkle et al., 2004; Tinkle et al., 2008). However, we find that increased levels of P-cadherin in keratinocytes does not result in compensatory upregulation of P-cadherin in melanoblasts of E-Cad KC cKO skin (Figure 5B). Thus, although melanoblasts in this model continue to express E-cadherin, there is little to no available E-cadherin on surfaces of surrounding keratinocytes to bind.

**Figure 5.**
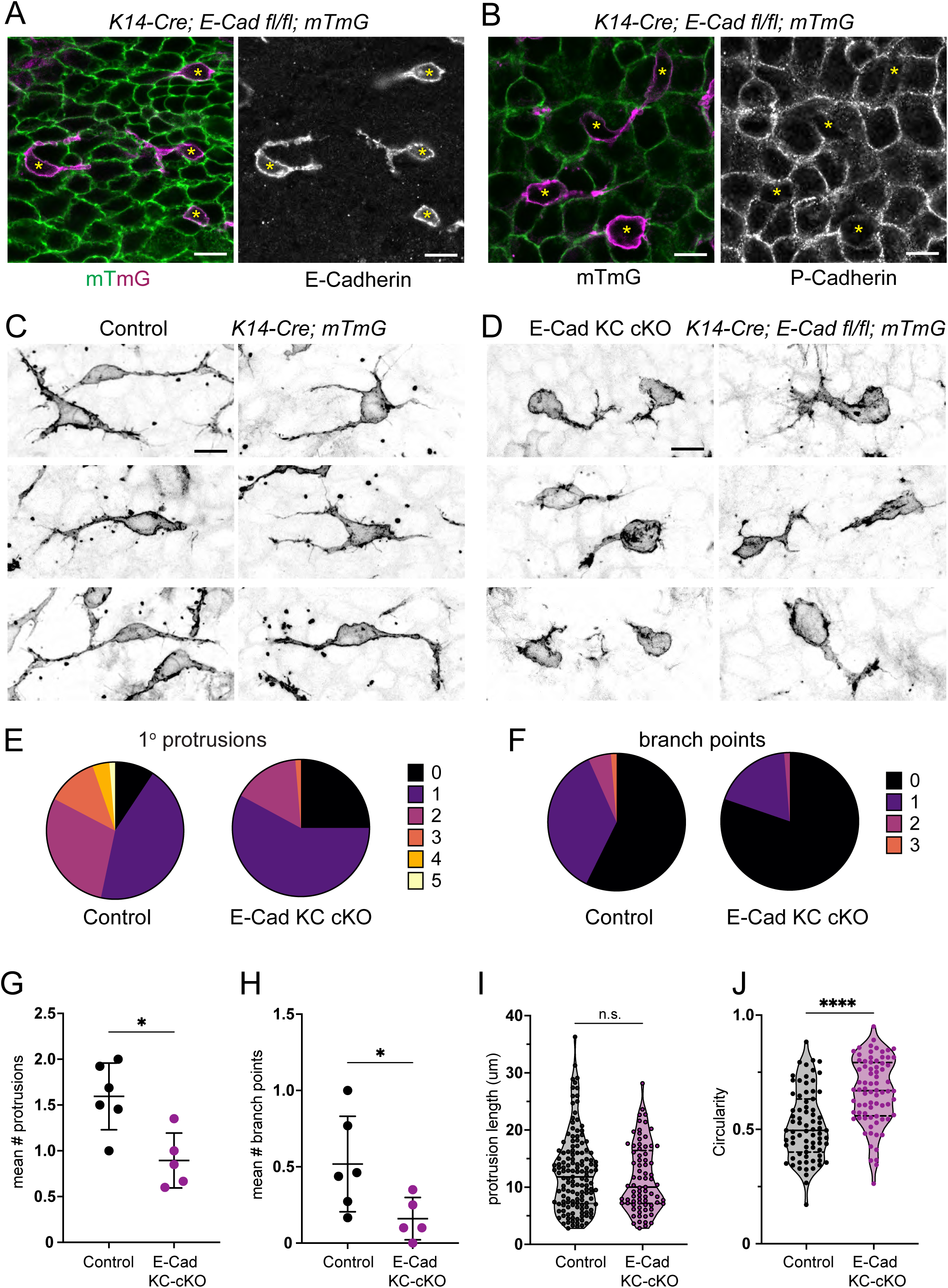
Melanoblasts require E-cadherin non-autonomously in surrounding keratinocytes to form stable migratory protrusions. **(A)** Representative planar views of wholemount skin explants from E16.5 *K14-Cre; E-Cad fl/fl; mTmG* embryos labeled with E-cadherin (right panel, grayscale). E-cadherin is selectively depleted in keratinocytes (green, mG) but not melanoblasts (magenta, mT). Yellow asterisks mark melanoblasts. Scale bar, 10um. **(B)** Representative planar views of wholemount skin explants from E16.5 *K14-Cre; E-Cad fl/fl; mTmG* embryos labeled with P-cadherin (right panel, grayscale). P-cadherin is upregulated in keratinocytes (green, mG) but not melanoblasts (magenta, mT), compare to Figure 2B. Yellow asterisks mark melanoblasts. Scale bar, 10um. **(C-D)** Representative melanoblasts from E16.5 control (*K14-Cre; mTmG,* C) and E-Cad KC cKO (*K14-Cre; E-Cad fl/fl;* mTmG, D) wholemount backskins. Shown are z-projections of the mT channel labeling melanoblasts in inverted grayscale to highlight melanoblast morphology. Scale bars, 10um. **(E-F)** Pie charts depicting the number of primary protrusions (E) and branch points per primary protrusion (F) displayed by control and E-Cad KC cKO melanoblasts. n=75 control and n=76 Ecad KC cKO melanoblasts. **(G)** Mean number of primary protrusions across melanoblasts per biological replicate. n=5 control and n=6 Ecad KC cKO embryos. Mann-Whitney test p=0.0108. **(H)** Mean branch points across melanoblasts per biological replicate. n=5 control and n=6 Ecad KC cKO embryos. Mann-Whitney test p=0.0281. **(I)** Length of primary protrusions in um. Mann-Whitney test p=0.4395 n.s. **(J)** Circularity ratio. Mann-Whitney test p<0.0001. For E-J, n=75 melanoblasts across 6 control embryos, and n=76 melanoblasts across 5 E-Cad KC cKO embryos. For G-J mean +/- S.D. are shown.

To investigate the consequences of depleting epithelial E-cadherin on melanoblast morphology, we acquired confocal z-stacks through the epidermal layers of E-Cad KC cKO embryonic skin explants and generated maximum intensity projections of the melanoblast-expressing mTomato channel, which allowed us to visualize the full dendritic morphology of wild-type melanoblasts in an E-cadherin deficient environment (Figure 5C-D). Consistent with our findings in E-Cad MB cKO skin, we observed a reduction in the number of long, primary protrusions and secondary branches in melanoblasts surrounded by epithelial cells lacking E-cadherin (Figure 5E-H). Moreover, melanoblasts in E-Cad KC cKO epidermis displayed shorter primary protrusions and increased cell body circularity (Figure 5I-J). Collectively, these data indicate that the ability of melanoblasts to form stable migratory protrusions depends on homotypic E-cadherin binding at melanoblast-keratinocyte interfaces.

### Melanoblasts form atypical E-cadherin-based attachments to neighboring keratinocytes

Typically, motile cells that express E-cadherin migrate as collectives, using E-cadherin homotypic attachments to connect neighboring cells and couple their cytoskeletal dynamics to move as a group (Friedl and Mayor, 2017; Gupta and Yap, 2021; Mayor and Etienne-Manneville, 2016). And yet, despite their E-cadherin expression, melanoblasts migrate as individuals, squeezing between keratinocytes connected by previously established homotypic E-cadherin junctions. Thus, we hypothesized that melanoblasts form unique E-cadherin-based attachments, distinct from those connecting keratinocytes, that enable melanoblasts to migrate individually through the epidermal layers. In epithelia, E-cadherin homotypic attachments assemble into mechanically-resistant adherens junctions by connecting to the actin cytoskeleton via intracellular catenins (Meng and Takeichi, 2009; Troyanovsky, 2023). We therefore compared the localization of α-catenin, β-catenin and p120-catenin at MB-KC versus KC-KC junctions. Whole mount skin explants from *Tyr-Cre; mTmG* embryos were labeled with antibodies against α-catenin, β-catenin or p120-catenin and catenin levels were quantified by segmenting melanoblast or keratinocyte cell edges and calculating catenin mean edge fluorescence intensities per cell. Strikingly, α-catenin levels at MB-KC junctions were significantly lower than those at KC-KC edges (Figure 6A-B). This is in contrast to the high levels of E-cadherin present at MB-KC interfaces. (Figure 2). Similarly, β-catenin (Figure 6C-D) and p120-catenin (Figure 6E-F) levels were lower at MB-KC interfaces compared to KC-KC junctions. To determine if catenin expression is downregulated in melanoblasts at the transcriptional level, we performed a bioinformatic analysis of cell-cell adhesion gene expression across existing bulk and single-cell RNA sequencing datasets (Jacob et al., 2023; Johnson et al., 2023; Sennett et al., 2015). Analysis of datasets from E14.5 and E15.5 stages of embryonic development acquired by different groups showed that levels of α-catenin, β-catenin and p120-catenin transcripts in melanoblasts are roughly equivalent to those of keratinocytes (Figure S2). Thus, it is likely that the paucity of catenins at MB-KC interfaces is due to post-transcriptional regulation of catenin and/or E-cadherin activity.

**Figure 6.**
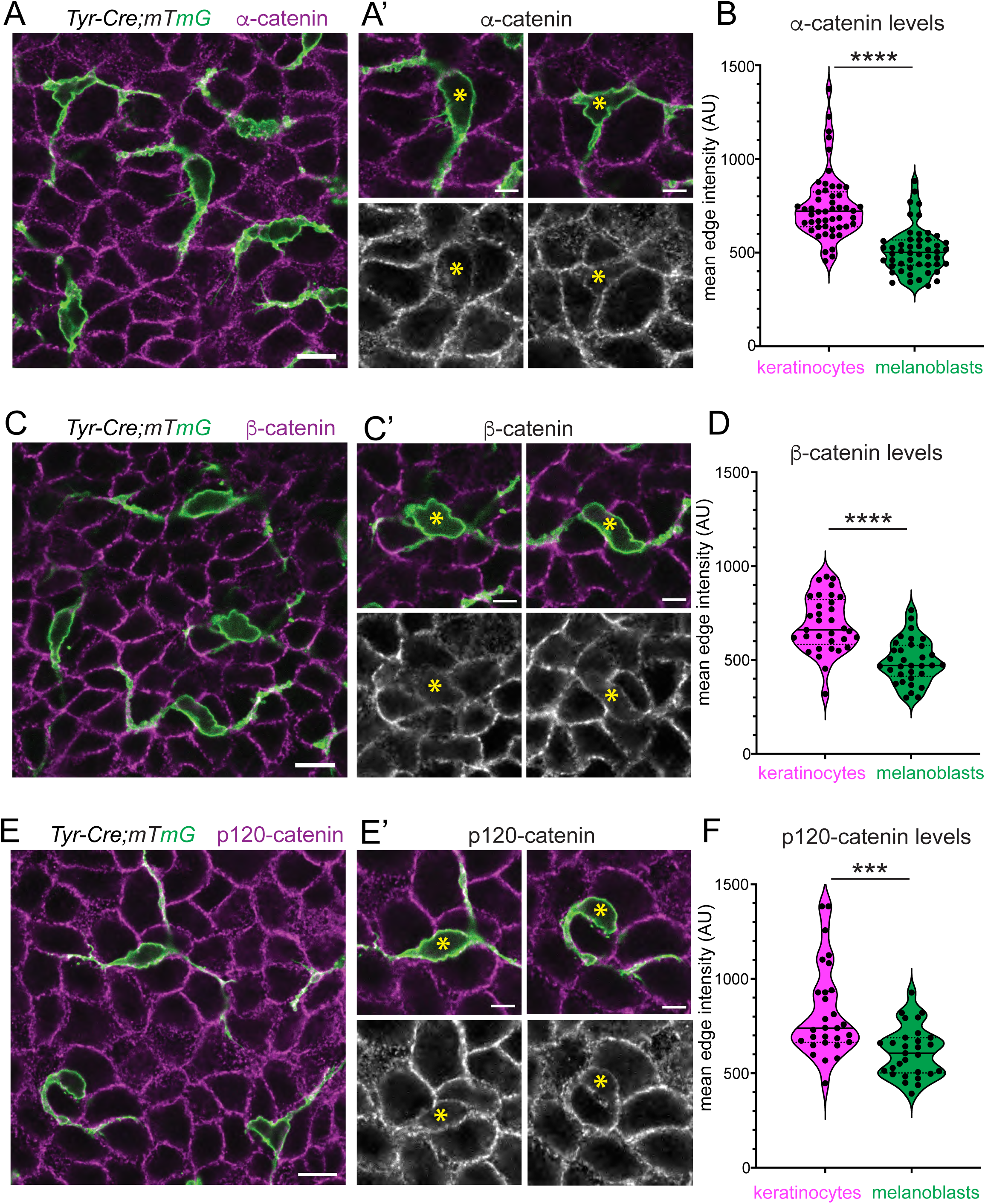
Melanoblasts form atypical E-cadherin-based attachments to surrounding keratinocytes that largely lack cytoplasmic catenins. **(A, C, E)** Representative planar views of wholemount skin explants from E16.5 *Tyr-Cre; mTmG* embryos labeled with α-catenin (A), β-catenin, (B) or p120-catenin (C) (magenta). Melanoblasts are labeled with mGFP (green). Scale bars, 10um. **(A’, C’, E’)** Two zoomed-in examples each of melanoblast-keratinocyte interfaces from their corresponding images in A, C and E. Lower panels show catenin channel alone (grayscale). Yellow asterisks mark melanoblast cell bodies. Scale bars, 5um. **(B, D, F)** Quantification of mean catenin intensity along KC-KC and MB-KC interfaces as indicated. Each point is the mean edge intensity per cell. n= 50 melanoblasts and 50 keratinocytes across three E16.5 skins (B, Mann-Whitney test p<0.0001), n=33 melanoblasts and 30 keratinocytes (D, Mann-Whitney test p<0.0001), and n=30 melanoblasts and 30 keratinocytes (F, Mann-Whitney test p=0.0001).

Together, these data demonstrate that melanoblasts form atypical E-cadherin-based attachments to neighboring keratinocytes that largely lack the intracellular catenins that connect to actin and are typically associated with stabilizing E-cadherin homotypic adhesions. This arrangement suggests that E-cadherin engagement with the actin cytoskeleton may be more transient in migratory melanoblasts compared to stationary epithelial cells, perhaps generating less rigid attachments that enable melanoblasts to adhere to, yet migrate through, the skin epithelium.

### E-cadherin localized to MB-KC junctions is more mobile than that at KC-KC junctions

Our findings thus far show that melanoblasts form extensive, yet atypical, E-cadherin based attachments to surrounding keratinocytes that are essential for melanoblast morphology and migration. Given the transient nature of melanoblast-keratinocyte interactions, and the low level of junctional catenins present at MB-KC interfaces, we hypothesized that E-cadherin may be more dynamic or turnover faster within the membranes of MB-KC junctions, allowing melanoblasts to move as individuals between epidermal layers. To compare E-cadherin dynamics at MB-KC versus KC-KC junctions, we performed Fluorescence Recovery After Photobleaching (FRAP) analysis of E-cadherin in epidermal explants expressing E-cadherin-CFP (Snippert et al., 2010). In this experiment, melanoblasts express cytoplasmic tdTomato driven by Tyr-Cre, and E-cadherin is visualized through a C-terminally fused CFP tag at its endogenous locus (*Tyr-Cre; E-cadherin-CFP; tdTomato*)(Madisen et al., 2010; Snippert et al., 2010). Thus, E-cadherin-CFP localized to MB-KC interfaces can be easily distinguished from that at KC-KC interfaces (Figure 7A-B). Regions along MB-KC or KC-KC junctions were selected for photobleaching and imaged continuously over a 2-minute recovery period (Figure 7A’-B’). E-Cadherin-CFP fluorescence at MB-KC junctions recovered faster and more completely compared to KC-KC junctions (Figure 7C). MB-KC interfacial E-Cadherin displayed a shorter half-life and larger mobile fraction than E-Cadherin-CFP at KC-KC interfaces (Figure 7D-E). Although these data do not distinguish whether the increased mobility of E-cadherin at MB-KC junctions is due to faster lateral diffusion within the membrane or to more rapid delivery via exocytosis/recycling, our results are consistent with a dynamic pool of E-cadherin-based attachments at the epidermal interfaces of migrating melanoblasts.

**Figure 7.**
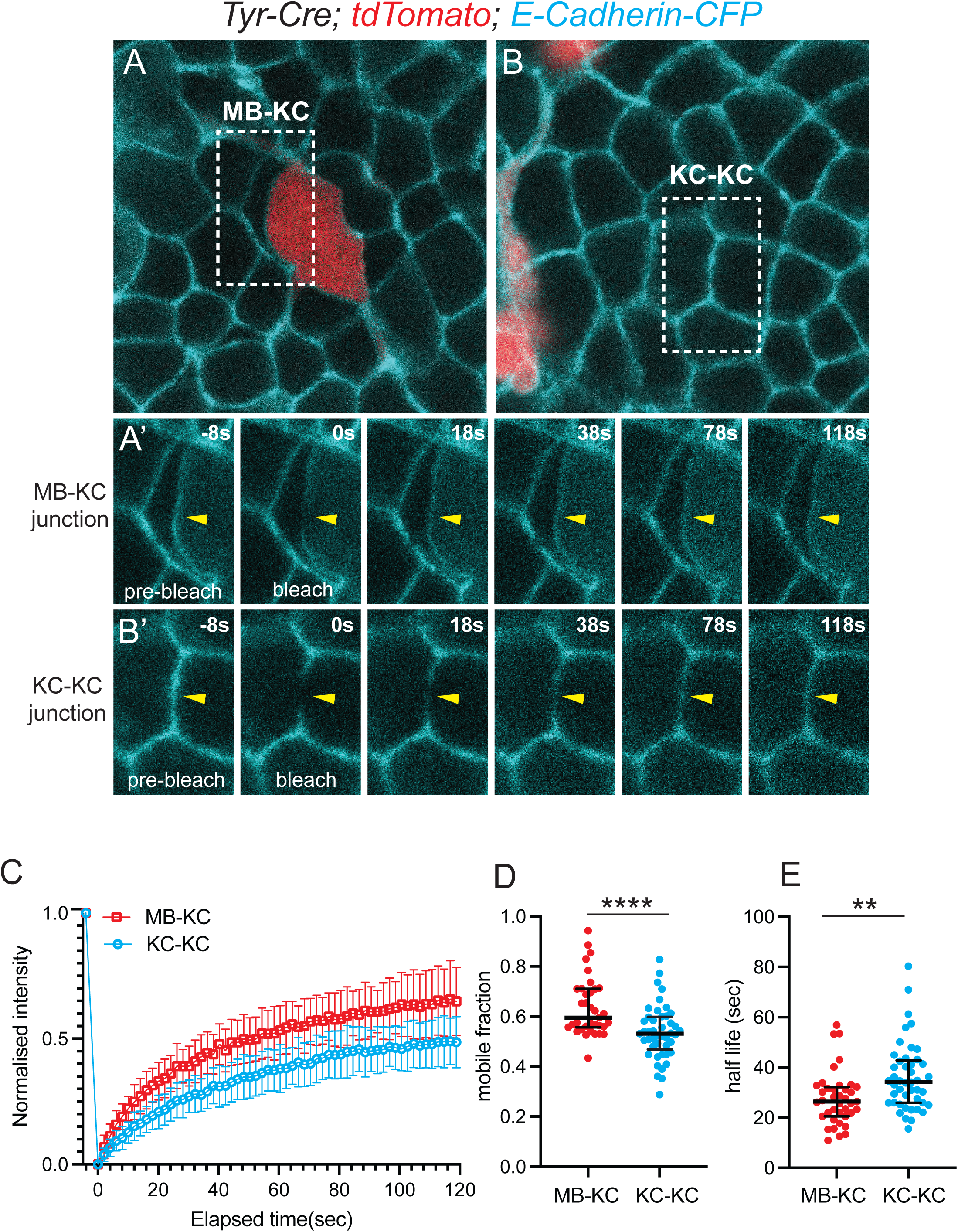
E-cadherin at melanocyte-keratinocyte interfaces is more mobile than keratinocyte junctions. **(A-B)** Representative still images from E16.5 *Tyr-Cre; tdTomato; E-cadherin-CFP* skin explants used from FRAP analysis of E-cadherin dynamics. tdTomato driven by Tyr-Cre marking melanoblasts (red) was used to identify E-cadherin (cyan) localized to melanoblast-keratinocyte (MB-KC) interfaces (A) versus keratinocyte-keratinocyte (KC-KC) junctions (B). White dotted lines mark FRAP regions shown below. **(A’-B’)** Representative still frames from FRAP analysis of E-cadherin-CFP at junctional regions shown in A (MB-KC) and B (KC-KC). 1 um regions of interest were photobleached and imaged at 2s intervals for 120s. Time points pre-and post-bleaching are indicated and time is shown in seconds (s). **(C)** E-cadherin-CFP fluorescence recovery curves at MB-KC and KC-KC interfaces as indicated. Data is pooled from n=44 traces across 3 embryos. Mean and S.D. are shown. **(D)** Mobile fraction of E-Cadherin-CFP at MB-KC vs KC-KC interfaces. Mann-Whitney test, p <0.0001. **(E)** Half-life of E-Cadherin-CFP at MB-KC vs KC-KC interfaces. Mann-Whitney test, p= 0.0012. For (D-E) each point represents one junction, bars display median and interquartile range.

## DISCUSSION

The mechanisms by which cells migrate upon and through extracellular matrices are well understood, but we know comparatively little about how accessory cells that populate epithelial tissues move through confined and adherent epithelial spaces (Pawluchin and Galic, 2022; Yamada and Sixt, 2019). Here we show that melanocyte precursors residing in the skin epidermis use, in addition to the basement membrane, epithelial surfaces as substrates for migration. Melanoblasts upregulate E-cadherin expression upon crossing into the epidermis (Nishimura et al., 1999), and here we have defined the form and function of E-cadherin-based adhesions in melanoblast migration. Melanoblasts form dynamic E-cadherin homotypic attachments to surrounding keratinocytes that are required to stabilize dendritic protrusions and promote . These attachments contain relatively little cytoplasmic catenins compared to the stable adhesions that form between epidermal cells, suggesting that melanoblasts employ distinct mechanisms to regulate junctional assembly and turnover. We propose that high levels of E-cadherin on the surfaces of melanoblast protrusions may outcompete homotypic E-cadherin interactions attaching neighboring keratinocytes, allowing melanoblasts to intercalate between them. However, because MB-KC attachments are not reinforced by appreciable levels of catenin recruitment and, presumably, F-actin binding, they remain relatively fluid within the membrane and display faster turnover than epithelial adhesions. This idea aligns with recent work demonstrating that actin-dependent α-catenin oligomerization contributes to clustering, stability and strength of adherens junctions (Troyanovsky et al., 2025). Interestingly, E-cadherin is not needed, either cell autonomously or in surrounding keratinocytes, for melanoblasts to extend migratory protrusions, suggesting the signals and machinery that induce actin polymerization at the leading edges are independent of E-cadherin engagement. It is likely other accessory cell types that colonize epithelial tissues, such as immune cells and peripheral neurons, employ similar E-cadherin based mechanisms to migrate in confined epithelial spaces.

The function of E-cadherin in melanoblast motility is distinct from its best understood role in cell migration, which is to make migration collective (Cheung and Horne-Badovinac, 2025; Friedl and Mayor, 2017; Mayor and Etienne-Manneville, 2016). Homotypic E-cadherin attachments physically link neighboring cells of the same type so they move as a group. E-cadherin attachments further promote collective movement by coupling actin dynamics across lateral junctions of neighbors, and by restricting front-facing protrusion and rear-facing contractions to the free edges of the cell collective (Cheung and Horne-Badovinac, 2025; Gupta and Yap, 2021; Mayor and Etienne-Manneville, 2016). Adhesion of melanoblasts to keratinocytes, by contrast, does not lead to coupling of their movement. Rather, melanoblasts slide between keratinocytes without visibly altering or disrupting keratinocyte morphology or behavior. Presumably, the low levels of cytoplasmic catenins present at MB-KC interfaces prevents E-cadherin-based attachments from making strong connections to the cytoskeleton, allowing melanoblasts to move through the epithelium as individuals. How the cytoplasmic catenins are differentially regulated at MB-KC versus KC-KC junctions is unclear, but regulation appears to occur post-transcriptionally as melanoblasts express α-catenin, β-catenin and p120-catenin transcripts at levels comparable to surrounding keratinocytes. Whether the catenins are selectively degraded or post-translationally modified in melanoblasts will be important to determine.

Several other cell types use E-cadherin to engage in heterotypic, cell-on-cell migration through different microenvironments. *Drosophila* border cells, for example, migrate on surrounding nurse cells in the ovary, where they use E-cadherin rather than the ECM to attach to and squeeze between nurse cells (Messer and McDonald, 2023; Mishra et al., 2019). Loss of E-cadherin expression in nurse cells alone impairs border cell directional migration, suggesting E-cadherin serves as a substrate for their migration (Cai et al., 2014; Fulga and Rorth, 2002; Niewiadomska et al., 1999). Cells within the border cell cluster also make homotypic E-cadherin contacts with each other to migrate as a collective, but what makes homotypic and heterotypic E-cadherin attachments distinct is not known (Cai et al., 2014; Niewiadomska et al., 1999). Primordial germ cells (PGCs) must migrate through various stromal and epithelial microenvironments as they transit to the somatic gonad, which requires dynamic regulation of cadherin-based adhesion (Barton et al., 2024; Schick et al., 2025). In zebrafish PGCs, E-cadherin mediates traction forces between individual PGCs and surrounding cells, and PGCs lacking E-cadherin function fail to confine retrograde flows that restrict bleb formation to the leading edge, impeding forward movement (Grimaldi et al., 2020; Kardash et al., 2010). Although the pseudopod-based migratory mode of melanoblasts is distinct from the bleb-based migration of PGCs, it will be interesting to determine whether E-cadherin coordinates actin assembly to focus pseudopod protrusivity and migration directionality.

Following melanoblast migration in embryonic skin, adhesion to keratinocytes continues to function as crucial component of the adult melanocyte microenvironment. In human skin, keratinocyte attachment controls melanocyte growth, differentiation, and pigment production and defects in E-Cadherin based attachment are associated with pigmentation defects and melanoma (Green et al., 2024; Wang et al., 2016). In vitiligo patients, for example, E-Cadherin is lost from the interfaces between melanocytes and keratinocytes (Wagner et al., 2015), and loss of E-cadherin is thought be a critical step in melanoma progression (Green et al., 2024; Kuphal and Bosserhoff, 2012; Tang et al., 1994). Melanoma cells typically switch from E-Cadherin to N-cadherin expression, which may allow them to escape keratinocyte control, interact with dermal fibroblasts and endothelial cells, and facilitate tumor spread (Hsu et al., 2000; Hsu et al., 1996; Sanders et al., 1999). We did not observe the escape of E-cadherin-deficient MBs into the dermis of embryonic skin, suggesting additional genetic changes are needed to for MBs to physically escape their epidermal microenvironment. Keratinocyte adhesion has also been shown to non-autonomously suppress melanoma in mice, where removal of the polarity protein Par-3 in keratinocytes promotes melanocyte proliferation, motility and melanoma progression non-cell autonomously through up-regulation of P-cadherin (Mescher et al., 2017). Deciphering mechanisms by which keratinocyte-melanocyte adhesion controls melanocyte behaviors like migration and proliferation will be important for understanding the transformation to metastatic disease.

## MATERIALS AND METHODS

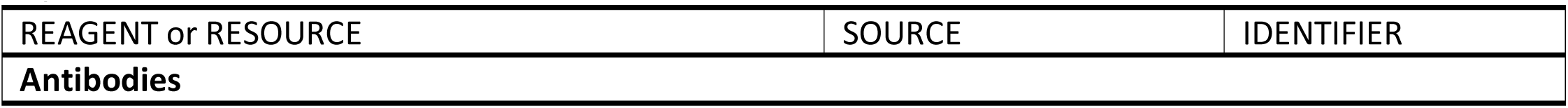
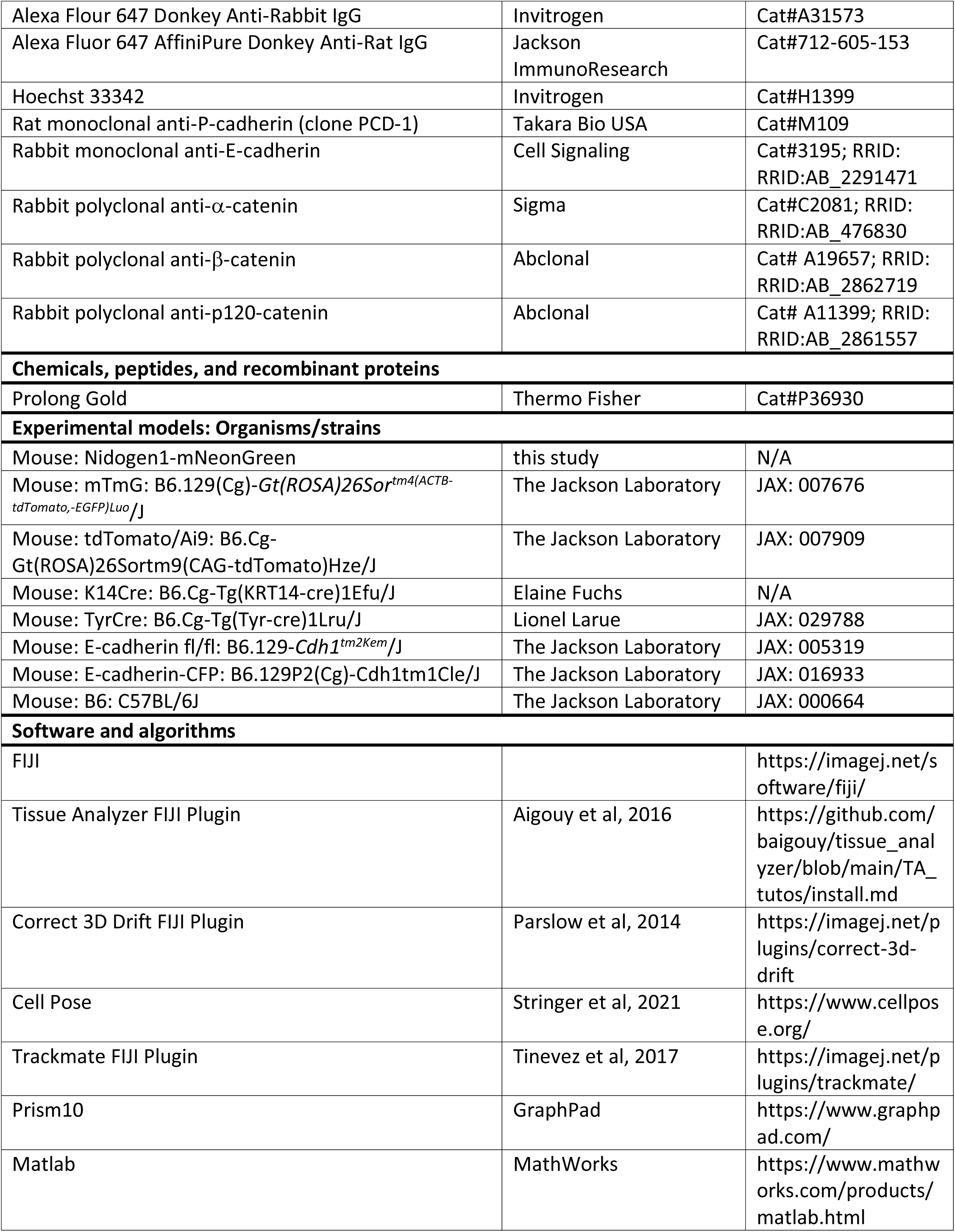
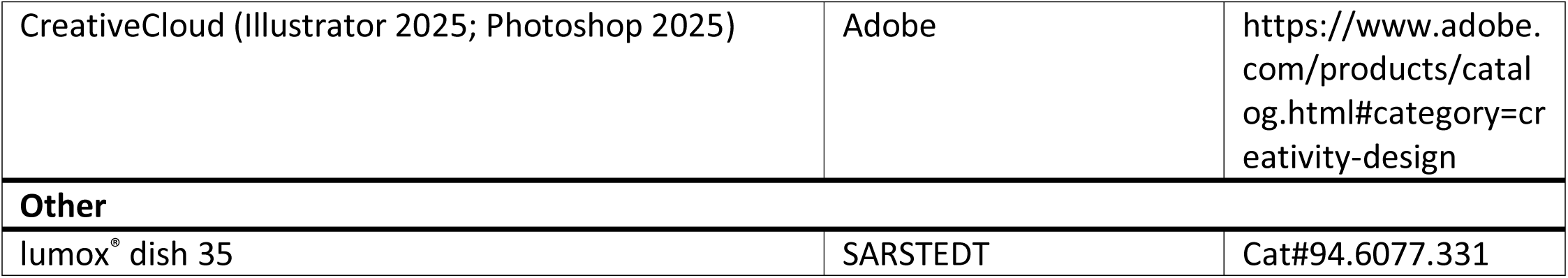
Key Resources Table

**Key Resources Table**
| REAGENT or RESOURCE | SOURCE | IDENTIFIER |
| --- | --- | --- |
| **Antibodies** |  |  |
| Alexa Flour 647 Donkey Anti-Rabbit IgG | Invitrogen | Cat#A31573 |
| Alexa Fluor 647 AffiniPure Donkey Anti-Rat IgG | Jackson ImmunoResearch | Cat#712-605-153 |
| Hoechst 33342 | Invitrogen | Cat#H1399 |
| Rat monoclonal anti-P-cadherin (clone PCD-1) | Takara Bio USA | Cat#M109 |
| Rabbit monoclonal anti-E-cadherin | Cell Signaling | Cat#3195; RRID: RRID:AB_2291471 |
| Rabbit polyclonal anti-α-catenin | Sigma | Cat#C2081; RRID: RRID:AB_476830 |
| Rabbit polyclonal anti-β-catenin | Abclonal | Cat# A19657; RRID: RRID:AB_2862719 |
| Rabbit polyclonal anti-p120-catenin | Abclonal | Cat# A11399; RRID: RRID:AB_2861557 |
| **Chemicals, peptides, and recombinant proteins** |  |  |
| Prolong Gold | Thermo Fisher | Cat#P36930 |
| **Experimental models: Organisms/strains** |  |  |
| Mouse: Nidogen1-mNeonGreen | this study | N/A |
| Mouse: mTmG: B6.129(Cg)-Gt(ROSA)26Sor<sup>tm4</sup>(ACTB-tdTomato,-EGFP)<sup>Luo</sup>/J | The Jackson Laboratory | JAX: 007676 |
| Mouse: tdTomato/Ai9: B6.Cg-Gt(ROSA)26Sortm9(CAG-tdTomato)Hze/J | The Jackson Laboratory | JAX: 007909 |
| Mouse: K14Cre: B6.Cg-Tg(KRT14-cre)1Efu/J | Elaine Fuchs | N/A |
| Mouse: TyrCre: B6.Cg-Tg(Tyr-cre)1Lru/J | Lionel Larue | JAX: 029788 |
| Mouse: E-cadherin fl/fl: B6.129-Cdh1<sup>tm2Kem</sup>/J | The Jackson Laboratory | JAX: 005319 |
| Mouse: E-cadherin-CFP: B6.129P2(Cg)-Cdh1tm1Cle/J | The Jackson Laboratory | JAX: 016933 |
| Mouse: B6: C57BL/6J | The Jackson Laboratory | JAX: 000664 |
| **Software and algorithms** |  |  |
| FIJI |  | https://imagej.net/software/fiji/ |
| Tissue Analyzer FIJI Plugin | Aigouy et al, 2016 | https://github.com/baigouy/tissue_analyzer/blob/main/TA_tutos/install.md |
| Correct 3D Drift FIJI Plugin | Parslow et al, 2014 | https://imagej.net/plugins/correct-3d-drift |
| Cell Pose | Stringer et al, 2021 | https://www.cellpose.org/ |
| Trackmate FIJI Plugin | Tinevez et al, 2017 | https://imagej.net/plugins/trackmate/ |
| Prism10 | GraphPad | https://www.graphpad.com/ |
| Matlab | MathWorks | https://www.mathworks.com/products/matlab.html |
| CreativeCloud (Illustrator 2025; Photoshop 2025) | Adobe | https://www.adobe.com/products/catalog.html#category=creativity-design |
| **Other** |  |  |
| lumox® dish 35 | SARSTEDT | Cat#94.6077.331 |

### Mouse lines and breeding

All procedures involving animals were approved by Princeton University’s Institutional Animal Care and Use Committee (IACUC). Mice were housed in an AAALAC-accredited facility in accordance with the Guide for the Care and Use of Laboratory Animals. This study was compliant with all relevant ethical regulations regarding animal research. E15.5 – E16.5 embryos from C57BL6/6J backgrounds were used unless otherwise indicated. Both sexes were used as sex was not determined in embryos. Genotypes were determined by PCR analysis of each allele. The presence of *mTmG* was validated by screening embryos for red and/or green fluorescence. E-Cadherin protein depletion was validated by staining skin explants with E-Cadherin antibodies. A list of mouse strains used in this study is provided in the Key Resources Table.

### Whole-mount immunofluorescence

For immunostaining, E15.5-E16.5 embryos were dissected in PBS and fixed in 4% paraformaldehyde at room temperature for 1 h. Backskins were dissected from fixed embryos and blocked for 1 h at room temperature or overnight at 4°C in 2% normal goat serum, 2% normal donkey serum, 1% bovine serum albumin and 1% fish gelatin in PBT3 (PBS with 0.3% Triton X-100). Skins were incubated with primary antibodies diluted in blocking solution overnight at 4°C. Skins were washed 3-5 times in PBT3 and incubated with secondary antibodies for 2 h at room temperature or overnight at 4°C in PBT3. When samples were stained using the P-Cadherin antibody, TBS with 0.3% Triton X-100 was used instead of PBT3 for all steps. Full thickness backskins were flat mounted in Prolong Gold. The following primary antibodies were used: rabbit anti-E-cadherin (1:500, Cell Signaling, Cat: 3195), rat anti P-cadherin (1:200, Invitrogen, Cat: 13-2000Z), rabbit anti α-catenin (1:2000, Sigma, Cat: C2081), rabbit anti β-catenin (1:500, Abclonal, Cat: A19657), rabbit anti p120-catenin (1:500, Abclonal, Cat: A11399). Alexa Fluor-647 secondary antibodies were used at 1:1000. Hoechst (Invitrogen, Cat: H1399) was used at 1:1000. Images were acquired on an inverted Nikon A1R-STED confocal microscope controlled by NIS Elements software using a Plan Apo 100x/1.45N.A. oil immersion objective.

ImageJ/FIJI was used for image processing.

### Live imaging

E16.5 dorsal skin explants were dissected in PBS and transferred to a 1% agarose gel supplemented with F-media (DMEM:F12 containing 10% fetal bovine serum). Explants were sandwiched between the gel on the dermal side and a 35-mm air permeable lummox membrane dish (Sarstedt) on the epidermal side. Z-series with a 1-micron step size were acquired at 3 or 10-minute intervals for 3 – 20 hrs. Images were acquired using a Nikon W1 using a Plan Apo 20x/0.75 NA air objective and 1× or 2.8x optical zoom. Explants were cultured in a humid imaging chamber at 37°C with 5% CO_2_ during the course of imaging.

### Quantification of melanoblast morphology

Max intensity z-projections of confocal z-stacks from fixed tissue explants were used to quantify melanoblast morphological characteristics. Cell body circularity was measured using FIJI/ImageJ by manually defining an ROI around the melanoblast perimeter. The number of primary protrusions and branch points were manually counted. Protrusion length was measured using the line drawing feature in FIJI/ImageJ.

### Quantification of junctional cadherin, catenin levels

Basal epidermal cells and melanoblast cell edges labeled with E-cadherin, P-cadherin, α-catenin, β-catenin or p120-catenin were segmented using Cellpose and hand corrected using the ImageJ plugin, Tissue Analyzer. The mean junctional intensity per cell was calculated by determining the average pixel intensity surrounding the predefined cell segmentation, as previously described (Leybova et al., 2024).

### Movie processing and cell tracking

Timelapse movies were corrected for drift using the ImageJ plugin, Correct 3D drift (Parslow et al., 2014). Melanoblasts and keratinocytes were segmented using Cellpose and manually corrected using Tissue Analyzer(Aigouy et al., 2016; Stringer et al., 2021). Cell trajectories were analyzed in 310um x 310um fields of view using the FIJI plugin Trackmate. The centroid position of cell bodies were tracked at 10-minute intervals over 14 hours (Tinevez et al., 2017).

### Fluorescence recovery after photobleaching (FRAP)

MB-KC junctions in *Tyr-Cre; Ecad CFP/CFP; tdTomato* embryonic explants were imaged using a Plan Apo 100x/1.45 NA oil immersion objective (Nikon) with appropriate optical zoom on an inverted Nikon A1R laser scanning confocal microscope equipped with a stage-top Tokai Hit incubation chamber to maintain 37°C and 5% CO_2_. 1 μm diameter circular ROIs were bleached using a 445 nm laser and recovery was monitored for a period of 2 mins with 2-second intervals from MB-KC junction in the IFE. Three reference pre-bleach images were also acquired. Magnification, laser power (both for bleach and acquisition), pixel dwell time and acquisition rate were kept uniform across all measurements. The acquired images and measured ROIs in the time series were checked for Z-drift and corrected for presence of any XY drift. A reference ROI was made in a non-bleached region to correct for overall bleaching during image acquisition. The ROI values were extracted from drift corrected images in NIS Elements software and subsequently processed in Microsoft Excel and GraphPad Prism. Each image time series was background/autofluorescence subtracted and bleach corrected (to be referred to as corrected intensity henceforth) and thereafter the corrected intensity profile was normalized as (F_t_ − F_bleach_)/ (F_ini_ − F_bleach_), where, F_t_ is the corrected intensity of the ROI at a given time point, F_bleach_ is the corrected intensity at the time point immediately after bleaching and F_ini_ is the mean ROI intensity of the three pre-bleach frames. Each mean recovery curve was fitted to exponential one phase association equation in GraphPad Prism with an r-squared value > 0.85 to determine the fitted plateau and Y0 values, which were then used to determine the mobile fraction = [(Plateau−Y_0_)/(1− Y_0_)]. Data represented is pooled from three biological replicates.

### Statistics and Reproducibility

Data between two groups were compared using a two-tailed, unpaired Student’s *t*-test. An F-test was performed to compare variances, and if significantly different, the t-test was performed with Welsh’s correction. Prism (GraphPad) was used for these analyses and to plot the data. See figure legends for specific p-values and n-values. P-values > 0.05 were considered not to be significant (n.s.). All experiments presented in this study were performed on epidermal explants from at least 3 embryos. Representative images have associated quantification and statistical analysis included in the legends.

## ACKNOWLEDGEMENTS

We are grateful to Gary Laevsky and Sha Wang of the Princeton Confocal Core Facility, a Nikon Center of Excellence for assistant with microscopy; Katie Little for assistance with mouse husbandry and genotyping; Rishabh Sharan for assistance with image analysis; members of the Devenport lab, Rebecca Burdine and Dan Notterman for helpful discussions. This research was supported by NIH NIAMS NRSA award F30 AR081690 (D.R.); NIH NICHD grant R01 HD105009 (D.D.); NIH NIAMS grant R01AR068320 (D.D), NIH NIGMS T32GM007388 (B.T.), the New Jersey Alliance for Clinical and Translational Science (D.D., R.J, B.T.), the Ludwig Institute for Cancer Research (D.D. and R.J), and the New Jersey Commission for Cancer Research (P.S).

## SUPPLEMENTAL FIGURE LEGENDS

**Figure S1.**
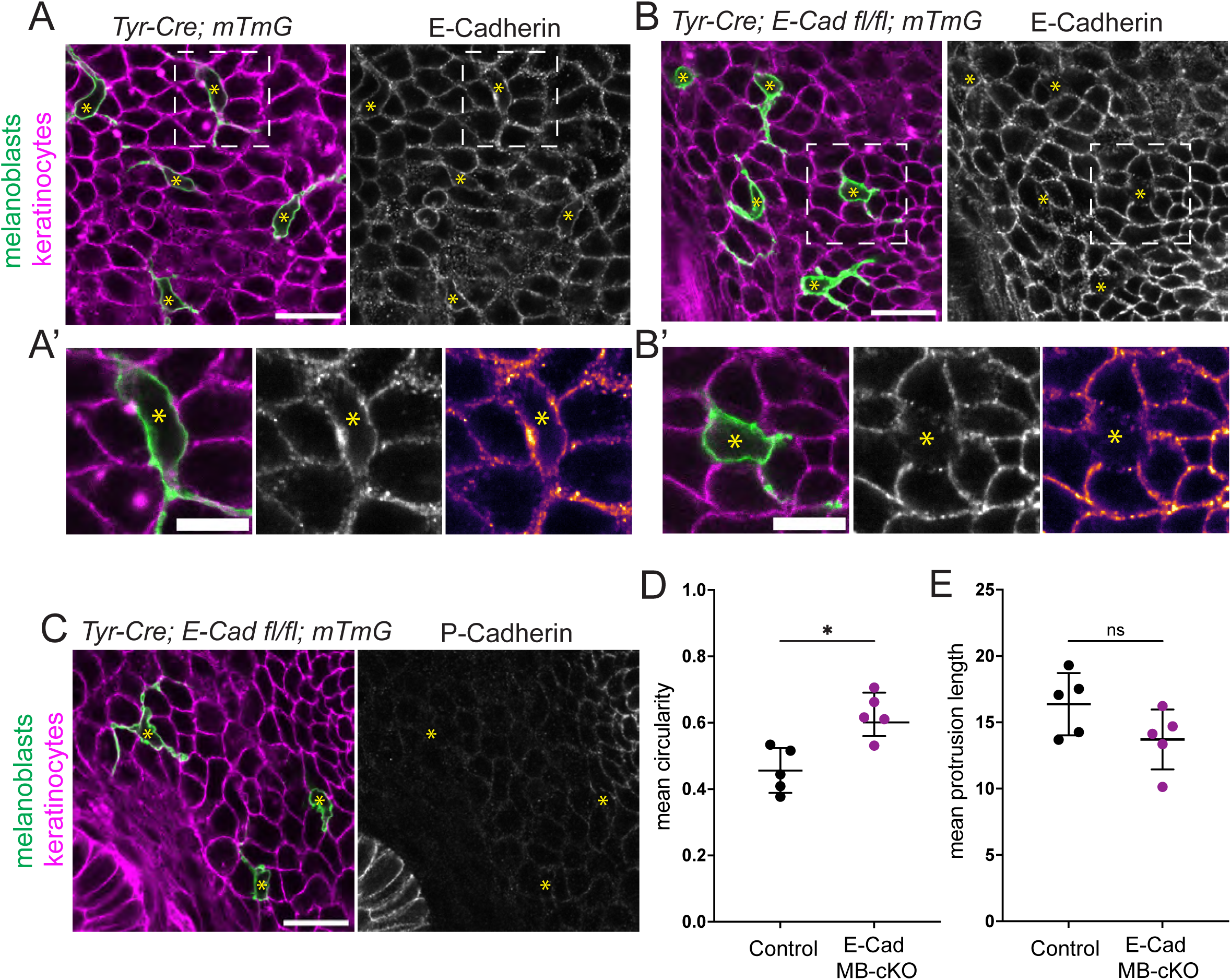
Depletion of E-cadherin in embryonic melanoblasts. **(A)** E16.5 *Tyr-Cre; mTmG* control whole mount skin explant labeled with E-cadherin. Melanoblasts are labeled in green (mG) and keratinocytes are in magenta (mT)(left panel). E-cadherin localizes at MB-KC interfaces (grayscale, right panel). **(A’)** Zoomed in view of melanoblast outlined by dashed box in A. Right panel shows E-Cad intensity LUT. Yellow asterisks mark melanoblast cell bodies. **(B)** E16.5 *Tyr-Cre; E-Cad fl/fl; mTmG* (MB E-Cad KO) mutant explants labeled for E-cadherin. **(B’)** Zoomed in view of melanoblast outlined by dashed box in B. Right panel shows E-Cad intensity LUT and confirms effective depletion of melanoblast-specific E-cadherin. Yellow asterisks mark melanoblast cell bodies. (**C**) E16.5 *Tyr-Cre; E-Cad fl/fl; mTmG* mutant explants (left) stained for P-cadherin (right, grayscale). P-cadherin is not upregulated to compensate for loss of E-cadherin. Yellow asterisks mark melanoblast cell bodies. **(D)** Mean cell body circularity per biological replicate. Average of 67 melanoblasts across n=5 control embryos and average of 72 E-Cad cKO melanoblasts across n=5 MB E-Cad cKO embryos. Mean +/-S.D. are shown. Mann-Whitney test. p=0.016*. **(E)** Mean protrusion length per biological replicate. Average of 67 melanoblasts across n=5 control embryos and 72 Ecad cKO melanoblasts across n=5 MB E-Cad cKO embryos. Mean +/- S.D. are shown. Mann-Whitney test. p=0.151 n.s.

**Figure S2.**
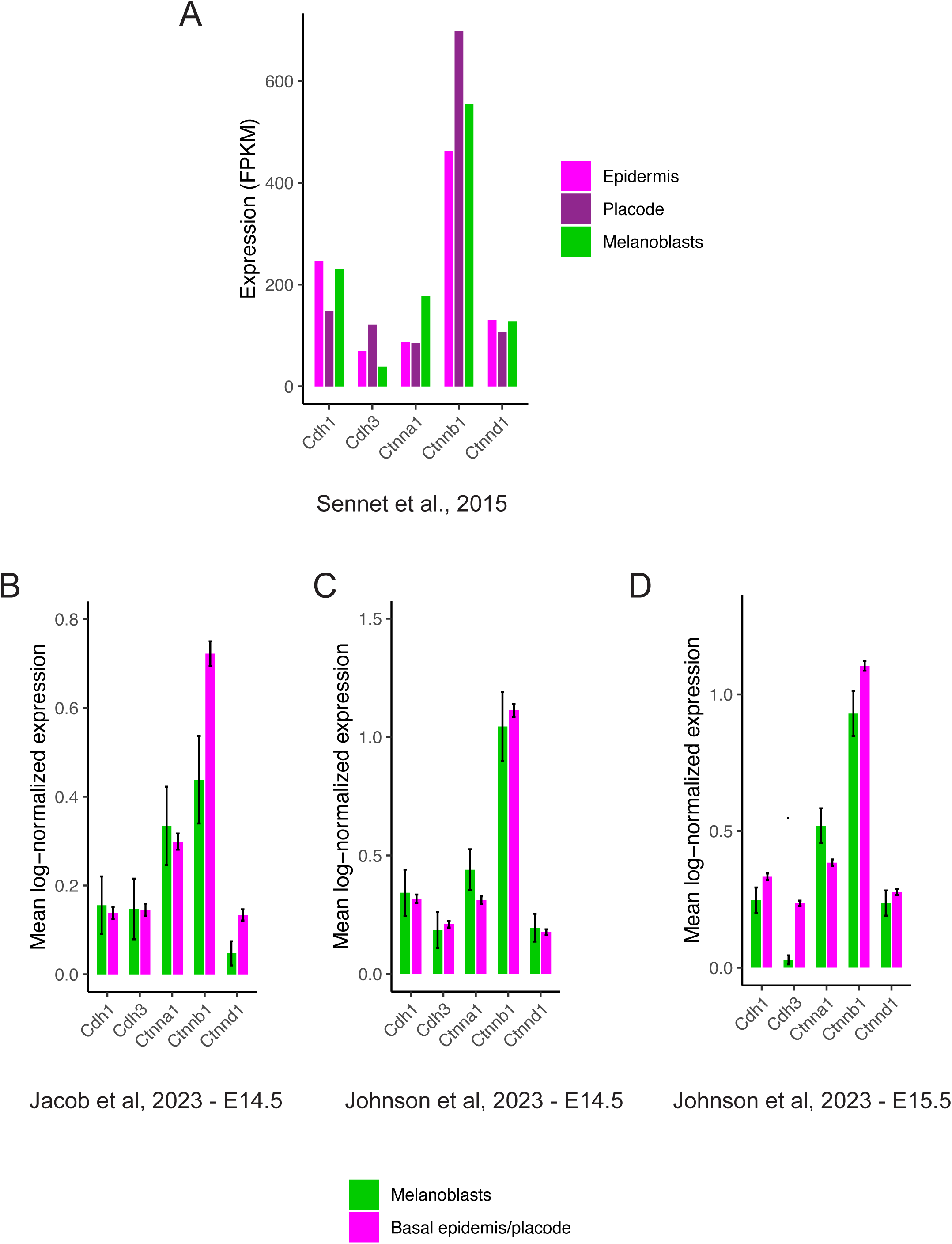
Transcriptional analysis of adherens junction components in melanoblasts versus keratinocytes. Analysis of Cdh1 (E-cadherin), Cdh3 (P-cadherin), Ctnna1 (α-catenin), Ctnnb1 (β-catenin), and Ctnnd1 (p120-catenin) transcripts from previously published bulk (Sennett et al., 2015) and single-cell (Jacob et al., 2023; Johnson et al., 2023) RNA sequencing datasets of embryonic skin. **(A)** Bulk transcript levels of indicated genes profiled from FACS sorted melanoblasts and interfollicular epidermal cells (IFE) (Sennett et al., 2015). **(B-D)** Analysis of adherens junction transcripts from ssRNA sequencing datasets of embryonic skin at E14.5 and E15.5 Melanoblasts are defined as expressing at least one of *Dct, Tyr, Pmel, Mlana.* Basal interfollicular epidermal plus hair follicle placode cells defined as expressing *Krt14* and/or *Krt5*. Bar plots show the mean log-normalized expression (± standard error) of the five genes. **(B)** Data from Jacobs et al, 2023. Embryonic skin at E14.5 n = 76 melanoblasts, n = 1882 basal epidermis/placode. **(C)** Data from Johnson et al, 2023. Embryonic skin at E14.5 n = 52 melanoblasts, n = 1209 basal epidermis/placode. **(D)** E15.5 data from Johnson et al, 2023. n = 126 melanoblasts, n = 2971 basal epidermis/placode. Statistical significance was assessed using the Wilcoxon rank-sum test with Bonferroni correction - no genes showed statistically significant differential expression between the two groups.

## SUPPLEMENTAL MOVIE LEGENDS

**Movie S1. Melanoblasts migrate between epidermal keratinocytes.** Time-lapse images of skin explant from E16.5 *Tyr-Cre; mTmG* embryo. mGFP-expressing melanoblasts (green) migrate between mTomato-expressing keratinocytes (magenta). Time is shown in hours and minutes. Scale bar, 10um.

**Movie S2. Melanoblasts migrate between keratinocytes of developing hair follicles.** Time-lapse images of skin explant from E16.5 *Tyr-Cre; mTmG embryo*. mGFP-expressing melanoblasts (green) migrate between mTomato-expressing keratinocytes of developing hair follicles (magenta). Time is shown in hours and minutes. Scale bar, 10um.

**Movie S3. Melanoblasts migrate along both basal and surpabasal layers.** PS Multiview imaging of skin explant from E16.5 *Tyr-Cre; tdTomato; Nid1-mNG embryo*. Sagittal view is shown. tdTomato-expressing melanoblasts (magenta) migrate along and near the basement membrane (BM) marked by Nid1-mNG (green), but also above the BM in the suprabasal layers of the epidermis. Time is shown in hours and minutes. Scale bar, 10um.

**Movie S4. Melanoblasts migrate without appreciable contact with the basement membrane in developing hair follicles.** PS Multiview imaging of skin explant from E16.5 *Tyr-Cre; tdTomato; Nid1-mNG* embryo. Sagittal view is shown. tdTomato-expressing melanoblasts (magenta) migrate along and near the basement membrane (BM) marked by Nid1-mNG (green), but also far from the BM in the center of the developing hair follicle. Time is shown in hours and minutes. Scale bar, 25um.

**Movie S5. Protrusion dynamics in wild-type control melanoblasts.** Time-lapse images of melanoblast from skin E16.5 *Tyr-Cre; mTmG* skin explant. mGFP channel is shown (grayscale). Time is in minutes. Scale bar, 10um.

**Movie S6. Protrusion dynamics in E-Cad MB cKO melanoblasts.** Time-lapse images of melanoblast from skin E16.5 *Tyr-Cre; E-Cad fl/fl; mTmG* skin explant. mGFP channel is shown (grayscale). Time is in minutes. Scale bar, 10um.

**Movie S7. Reduced protrusion stability in melanoblasts lacking E-cadherin.** Time-lapse images of melanoblast protrusions from E16.5 *Tyr-Cre; mTmG* (*control*, top) and *Tyr-Cre; E-Cad fl/fl; mTmG* (*E-Cad cKO*; bottom) skin explants. mGFP channel is shown (grayscale). Time is in minutes. Scale bar, 5um.

**Movie S8. Long-term imaging of melanoblast migration.** Time-lapse images of melanoblast migration in E16.5 *Tyr-Cre; mTmG* (*control*; left) and *Tyr-Cre; E-Cad fl/fl; mTmG* (*E-Cad cKO*; right) skin explants over 14 hours. mGFP channel is shown (inverted grayscale). Time is in minutes. Scale bar, 100 um.

